# Combined effects of mechanical loading and Piezo1 chemical activation on 22-months-old female mouse bone adaptation

**DOI:** 10.1101/2025.02.05.636720

**Authors:** Quentin A. Meslier, Robert Oehrlein, Sandra J. Shefelbine

**Author notes:** <u>Disclosures:</u> The authors declare no conflicts of interest.

## Abstract

With age, bones mechanosensitivity is reduced, which limits their ability to adapt to loading. The exact mechanism leading to this loss of mechanosensitvity is still unclear, making developing effective treatment challenging. Current treatments mostly focus on preventing bone mass loss (such as bisphosphonates) or promoting bone formation (such as Sclerostin inhibitors) to limit the decline of bones mass. However, treatments do not target the cause of bone mass loss which may be, in part, due to the bone’s inability to initiate a normal bone mechanapdatation response. In this work, we investigated the effects of 2 weeks of tibia loading, and Piezo1 agonist injection *in vivo* on 22-month-old mouse bone adaptation response. We used an optimized loading profile, which induced high fluid flow velocity and low strain magnitude in adult mouse tibia. We found that tibia loading and Yoda2 injection have an additive effect on increasing cortical bone parameters in 22-month-old mice. This combination of mechanical and chemical stimulation could be a promising treatment strategy to help promote bone formation in patients who have low bone mass due to aging.

## 1. Introduction

With age, the bone remodeling process is altered. Bone resorption exceeds the formation leading to a thinner and more porous tissue. This unbalance leads to higher risk of fracture burdening patients and the healthcare system [1,2]. Osteocytes are also affected by aging as their density, in the cortical bone, decreases with age [3]. The lacunar canalicular network (LCN), which connects osteocytes together, is disrupted in old mice bone [3,4]. Disruption of the LCN network could make the diffusion of key molecules challenging the osteocytes as well as the propagation of mechanical stimuli (e.g. fluid flow). Osteocytes have also been shown to express more senescence markers with aging [5–7], which could participate in the age-related bone mass loss [7]. These changes in bone structure and cellular activity most likely contribute to the changes in mechanoadpatation with aging.

Mechanical stimulation that typically induce bone adaptation in adult bone has shown disappointing results in older subjects [8–11]. This loss in bone mechanoresponsive with aging could be explained by the altered osteocytes capacity to sense and transduce the mechanical signal properly. Indeed, the number of osteocytes responding to mechanical stimulation in old mouse bone has been suggested to decrease compared to adult mice [12]. Studies showed that old mouse bone exhibited less upregulation of bone formation-related genes after loading compared to adult mouse bone [13–15], which could be explained, in part, by the disruption of the lacunar-canalicular network with aging [3,4]. In addition, bone mechanical environment has been shown to change under uniaxial compression load with aging [16]. Javaheri et al. [17] suggested that old mouse cortical bone adaptation can be primed by applying bouts of high magnitude load, which induced double the strain magnitude typically necessary for inducing bone formation in adult bone. These results suggested that commonly used loading profiles are insufficient to trigger a mechanoadaptation response in old mouse bone or suggest a dysfunction of the osteocytes mechanotransduction process at the commonly used load magnitude. Other approaches to restore old bone mechanoresponse included pharmaceutical treatments in combination with mechanical stimulation [18] [19].

Because the age-related bone mass loss is partially due to a loss of mechanosensitive processes in bone cells, finding ways to better stimulate osteocytes in old subjects could help in the design of new treatment strategy for patients with bone mass loss issue due to aging.

Osteocytes possess multiple mechanosensors to detect mechanical signals. Piezo1 is one of the main mechanosensitive ion channels in osteocytes and plays a critical role in bone mechanoadaptation response [20]. Piezo1 is required for bone development in embryos [21]. *In vitro*, Piezo1 mRNA expression and activity increased following fluid flow stimulation [20]. In mice, Piezo1 conditional knockouts in bone showed a decrease in cortical thickness and trabecular bone volume with the deletion of Piezo1 [20,22], and a diminished adaptation response to loading. In addition, mice placed in a situation of disuse via hindlimb unloading exhibited a decrease in expression of Piezo1 mRNA [22].

Yoda1 is a Piezo1 agonist that has been used to chemically stimulate bone cells. Yoda1 triggers calcium influx in bone cell cultures [20,22], which suggests an activation of the ion channels. This increase in calcium concentration was not observed in cells lacking expression of Piezo1. *In vivo*, 2 weeks of Yoda1 injections resulted in an increased bone mass, and an increase in bone formation-related mechanosensitive gene expression, such as *Wnt1*, in adult mice [20]. The hindlimb unloading model showed that Yoda1 injections mitigate bone loss induced by disuse in adult mouse bones [23].

As bones lose their ability to adapt with aging, Piezo1 agonists such as Yoda1 could help activate osteocytes and prevent bone mass loss. Recent studies investigated the effects of 4 weeks of Yoda1 injection in estrogen-deficient mice (ovariectomized at 8 weeks old) [23]. Yoda1 treatment slightly rescued bone loss induced by estrogen deficiency. In addition, this study showed a small but significant protective effect of Yoda1 injections in 20-months-old mice trabecular bone but not cortical bone parameters, suggesting that Yoda1 injection might participate in reducing bone mass loss[23]. Recently, Wasi et al. [24] investigated the combined effects of 4 weeks of Yoda1 injections and moderate tibia loading (that typically does not induce adaptation on its own) in 50-weeks-old mice. The authors concluded that loading and Yoda1 injection could help mitigate aging-associated bone loss.

In 2023, a new Piezo1 agonist, Yoda2, was developed. Yoda2 is more soluble in aqueous solution and more efficient at activating Piezo1 channel than Yoda1 [25]. This improved Piezo1 agonist could more easily diffuse in the bloodstream via its enhanced solubility and could participate in greater cells activation following *in vivo* injections.

Altogether, previous results suggest that chemically stimulating Piezo1 could be a promising treatment strategy to prevent bone mass loss related to aging. However, it is still unclear if it can help to promote bone adaptation in old mouse bone, which already has poor bone mass and degraded bone structure. In this work, we investigate the combined effects of 2 weeks of Yoda2 and *in vivo* tibia loading in 22-month-old mice. In addition, we used *in vivo* two-photon imaging transgenic mouse model to investigate the activation of osteocytes following Yoda2 injections. We reported a combined effect of mechanical and chemical stimulation in old mouse bone adaptation, particularly an increase in cortical bone parameters.

## 2. Methods

### 2.1. Animals

Animal experiments presented in this work were performed with the approval of Northeastern University’s Institutional Animal Care and Use Committee (IACUC). For loading experiments, 22-month-old female C57BL/6J mice (n=20) were obtained from the National Institute on Aging.

For *in vivo* calcium signaling imaging, transgenic mice expressing the fluorescent calcium indicator protein GCaMP3 specifically in osteocytes were generated, as previously described [26], by mating Ai38 (B6;129S-Gt(ROSA)26Sortm38(CAG-GCaMP3)Hze/J) and *Dmp1*-Cre mice (B6N.FVB-Tg(Dmp1-cre)1Jqfe/BwdJ) expressing the Cre recombinase under the control of *Dmp1*, which is mostly expressed in osteocytes in bone. Mice expressing both GCaMP3 and DMP1-Cre genes were aged until they reached 20 to 22 months old (n=2).

All mice used in this study were kept in groups of 5 maximum, with 12-hour light and dark cycles, and fed with a regular diet. A total of 22 mice were involved in this study.

### 2.2. *In vivo* calcium signaling imaging

Transgenic mice were anesthetized using 2% to 3% isoflurane and kept on a heating pad. The left leg was shaved using depilatory cream. The skin of the anterior tibia was surgically opened and the tibialis anterior was removed to expose the tibia. The leg was stabilized using a custom plate inspired by a cranial window fixture for *in vivo* imaging of the brain. The leg was glued to the stabilizing plate which was then screwed down to a custom-made platform (Supplemental Figure 1). The sealant was carefully applied at the leg-plate interface to prevent leaks. Ultrasound gel mixed with Hank’s Buffer Salt Solution, containing calcium ions, was prepared and added on top of the tibia to allow immersion of the objective. Fluorescent signal from calcium ions in osteocytes was recorded *in vivo* using two-photon microscopy. Imaging was performed with an excitation wavelength of 920 nm, a 500-550 nm filter, a 20x magnification water immersion objective, and at a rate of 3 frames per second. The region of interest was located about ∼20 µm below the surface of the bone and imaged for at least 5 minutes.

Baseline fluorescent signal fluctuation was recorded in transgenic mice prior to Yoda2 injection (n=2) in two different ROIs per mouse. Then a mouse was injected intraperitoneally with Yoda2 *in vivo* (n=1).

Cells were manually tracked in ImageJ and fluorescence intensities were analyzed as a function of time. The first 30 seconds of each cell trace were used to calculate the cell signal standard deviation. Signal was normalized, detrended, and smoothed using MATLAB and previously described method [27]. A cell was defined as responding if its fluorescence signal exhibited a peak with a fluorescence intensity at least 2 times larger than the standard deviation of the signal.

### 2.3. *In vivo* mouse tibia loading

We used the uniaxial tibia compression model [28] and applied loading on the right tibia of 22-month-old mice (n=20). During loading experiments, mice were anesthetized via 2% isoflurane and maintained on a heating pad. The right knee and the ankle were positioned in custom-made cups to ensure the axial orientation of the tibia. The loading profile was previously designed using FEM [29] to induce high fluid flow velocity but low strain magnitude in adult mouse bone. Cyclic load was applied to 7N at 175 N/s for 100 cycles and a rest period of 1 second was inserted between each cycle. Loading bouts were performed 3 days a week for 2 weeks. The left legs were kept as internal controls.

### 2.4. *In vivo* Piezo1 agonist injection

Yoda2 (KC289, hellobio, USA) was resuspended in 1ml of 100% DMSO at a concentration of 23 mM and stored at −20°c. Yoda2 working solution was prepared in 0.9% (w/v) Sodium Chloride, 5% DMSO, and 10% (w/v) cyclodextrin SBE-B-CD, to facilitate solubilization of the drug. The saline solution was first sterilized via 0.2 µm sterile filter before adding the Yoda2 stock solution. 200 µl of working solution were injected intraperitoneally *in vivo* at 5μmol/kg. Vehicle treatment was prepared in saline solution (0.9% (w/v) Sodium Chloride) with 5% DMSO without adding Yoda2.

Mice were injected *in vivo* 5 days a week for 2 weeks (n=10). The side of the injection was alternated every day. When the right tibia of the mouse was loaded, injections were made on the left side.

### 2.5. MicroCT scanning and analysis

After 2 weeks of *in vivo* loading and injections, mice were sacrificed via CO_2_ inhalation and cervical dislocation. Loaded and controlled legs were collected, muscles and other external tissues were removed from the bone, and bones were stored at −20°C in a gauze soaked with 1xPBS. Samples were scanned using microcomputed tomography (10μm resolution, 70 kVp, 114 mA, 400 ms integration time) and scans were analyzed in ImageJ, using the BoneJ plugin [30], to determine bone parameters.

In each scan, the tibial diaphysis was manually segmented. Proximal and distal tibia-fibula junctions were used to define the start and the end of the region to analyze. Once the scan was binarized, the plugin BoneJ [30] was used to calculate cortical bone parameters along the bone length: cortical area (Ct.Ar), cortical thickness (Ct.th), minimum moment of area (Imin), maximum moment of area (Imax), marrow area (Ma.Ar), total area, (Tt.Ar), and cortical bone fraction (Ct.Ar/Tt.Ar) [31]. We also quantify the gross cortical bone porosity at 15% and 37% of the bone length. The gross porosity was calculated by considering the ratio of the measured cortical area (Ct.Ar) and the theoretical cortical area (Tt.Ar - Ma.Ar), as described the following formula : 1 – (Ct.Ar/(Tt.Ar – Ma.Ar)); Supplemental Figure 2 illustrates the calculation of the gross cortical porosity.

For trabecular bone analysis, the region to be analyzed was defined as a 1mm region located 180 µm distal to the proximal tibial growth plate. Using BoneJ, parameters such as trabecular thickness (Tb.Th), trabecular number (Tb.N), trabecular spacing (Tb.Sp), and bone volume fraction (BV/TV) were quantified.

### 2.6. Statistical analysis

Statistical analysis of cortical bone parameters was performed using non-parametric 1D Statistical Parametric Mapping (SnPM1D) [32], in MATLAB (open source package: spm1d.org, © T. Pataky) SnPM1D allowed us to run a continuous statistical analysis on 1D data sets along the tibia length. In this study, the data sets correspond to cortical bone parameters measured from individual each microCT scan’s slice along the tibia. SnPM1D uses measurement of individual slices along the bone as input and allows comparison along the entire length of the bone. Evaluation of the data set distribution showed that some parameters along the bone length were not normally distributed. For this reason, non-parametric inference was used in the statistical analysis [33]. The null hypothesis assumed that the difference in mean cortical bone parameter (Ct.Ar, Ct.Th, Imax, and Imin) between the two tested groups equals zero. In addition, we tested the relative change, in cortical bone parameters, between loaded and control legs, from the vehicle and Yoda2 groups. If the relative changes were positive and significantly different from zero, then the bone parameters in the loaded leg were significantly higher than in the control leg. If the relative changes were negative, we concluded that the bone parameters in the loaded leg were significantly lower than in the control leg, with a significance level of α=0.05. SnPM1D test results were presented in bar plots along the tibia midshaft for each experimental condition (Control + Vehicle: n=10 bones; Control + Yoda2 n=10 bones; Loaded + Vehicle: n=8 bones; Loaded + Yoda2: n=8 bones). The shaded regions of the bar plots indicate that the null hypothesis was rejected at a Type I rate of α= 0.05, at the specified location. Using SnPM tests allowed us to report with transparency the changes in bone parameters between the loading leg and control leg along the bone length.

Note: SPM1D multicomparison and 2-way ANOVA are not validated. To test statistical differences between the groups we ran paired SnPM1D, using a non-parametric equivalent of two-sample ttest, long the bone length.

In literature, statistical analysis is commonly performed to compare bone parameters at a specific location of the bone (e.g., 50% of the mid-shaft length). However, this process prevents the detection of statistical differences along the rest of the bone. For comparison with the broader literature, we analyzed Ct.Ar, Ct.Th, and Imin at 15%, 25%, 35% and 50% of the tibia length. These regions of the bone have been reported to present adaptation following tibia loading [34–36]. In addition, we measured the marrow area (Ma.Ar), total area (Tt.Ar), and the cortical bone fraction (Ct.Ar/Tt.Ar) at 15% and 50% of the bone length. We used unbalanced 2-way ANOVA to determine the significant effects of the two independent factors: loading and Yoda2 injection, as well as the interactions of those two factors on cortical bone measurements. The ANOVA was unbalanced due to the exclusion of two mice in the Yoda2 group. MicroCT scans showed signs of a broken left tibia (control) in one of the mice, the second had an abnormal amount of trabecular bone in both loaded and control legs. Post-hoc tests were performed using the Tukey-Kramer method to determine which pairs of group means are significantly different. In this work, we considered data sets as independent groups. We tested the normal distribution of the data sets for each analyzed location and found that they were normally distributed. The porosity data was not normally distributed. For this reason, we used a Kruskal–Wallis test, a non-parametric equivalent of one-way ANOVA to test for significant different between groups. Post-hoc tests were performed using the Tukey-Kramer method to determine which pairs of group means are significantly different.

For the trabecular bone statistical analysis, we tested the normal distribution of the data and used unbalanced 2-way ANOVA to test the individual effects and interaction effects of loading and Yoda2 injection. Post-hoc was performed using the Tukey-Kramer method to determine which pairs of group means are significantly different.

## 3. Results

### 3.1. *In vivo* calcium imaging

We investigated the potential efficacy of Yoda2 injection to chemically trigger calcium ions influx in osteocytes *in vivo*. We used transgenic mice expressing GCaMP3 under the control of an osteocyte’s specific promoter, *Dmp1*. GCaMP expression in osteocytes results in the fluorescence of calcium ions once in the cells. *In vivo* two-photon microscopy (Figure 1.A) was used to track the fluorescence intensity over time (Figure 1.B). We assumed a correlation between fluorescence intensity fluctuation and calcium ion concentration [26]. Figure 1.C shows the percentage of responding cells before and after Yoda2 injections for one transgenic mouse. 60 min after Yoda2 injections we measured an increase in the percentage of responding osteocytes following Yoda2 injection *in vivo* (∼45% responding cells after 60min) compared to baseline (3% responding cells on average) (Figure 1.C). Baseline measurements were consistent between mice (n=2) with a percentage of responding cells fluctuating between 0% and 8%. Cells responding toYoda2 injection after 60 min exhibited peaks of fluorescence intensity with an average width of about 37 seconds (Figure 1.D), which matches previous results found in *ex vivo* mouse tibia [12,37].

**Figure 1:**
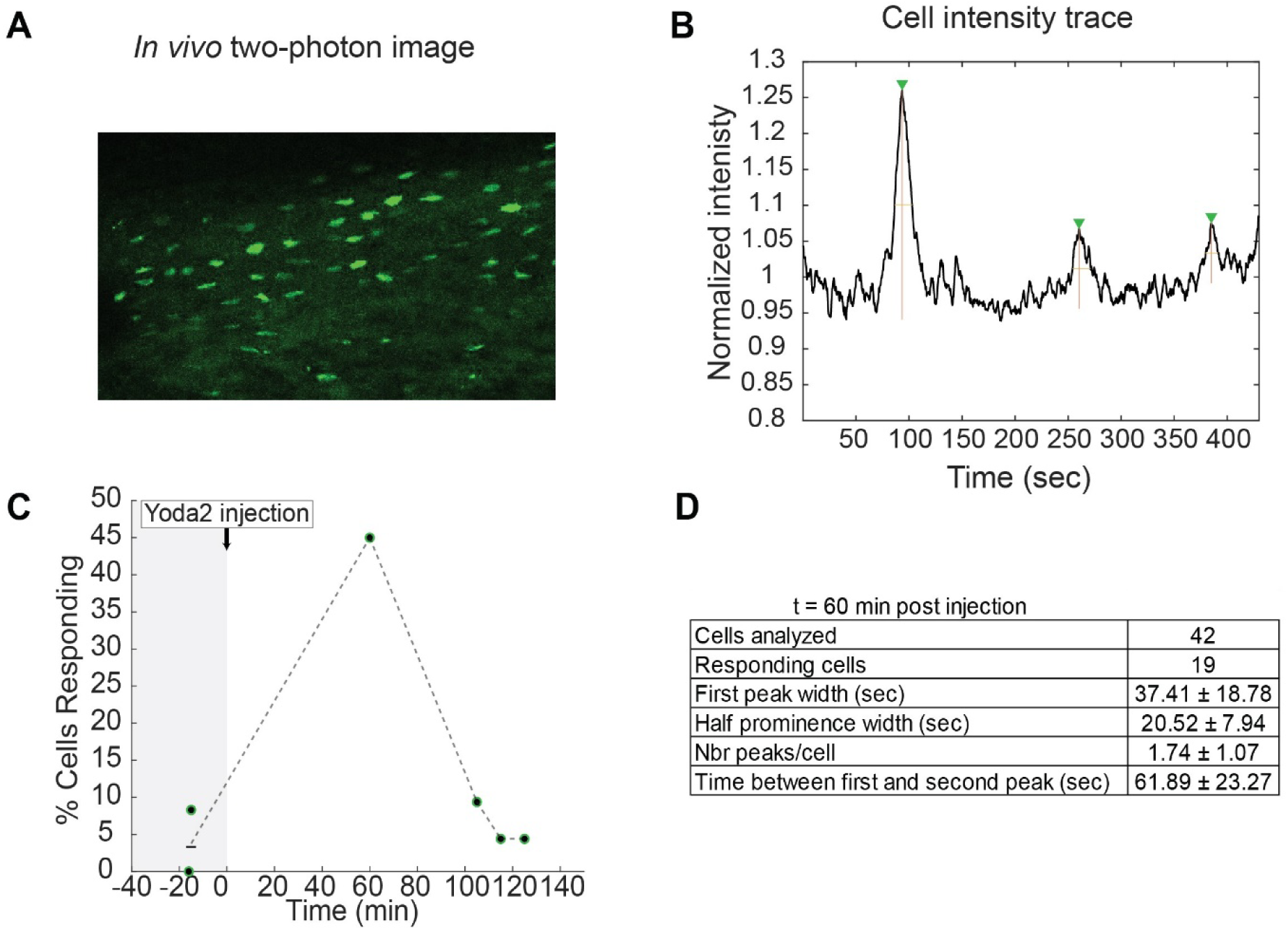
Quantification of in vivo calcium signaling in osteocytes in 22-month-old mice.In vivo two-photon microscopy image showing fluorescent osteocytes in the tibia of transgenic mice. B) Example of cell intensity trace over time after Yoda2 injection. Fluorescence intensity is associated with calcium concentration in osteocytes. The intensity trace shows fluorescent peaks interpreted as calcium signaling in flux. C) Percentage of responding cells at different time points before and after Yoda2 injection. Injection time is noted as t=0min. Two different ROIs from the same mouse were imaged before injection D) Quantification of responding cells characteristics at t=60min after injection (n=1).

The number of responding cells was measured to be similar to baseline approximately 2h after injection (Figure 1.D).

These results hint that Yoda2 was potentially able to reach osteocytes and trigger an increase in calcium concentration 60 min after *in vivo* injection which is consistent with previous studies looking at the diffusion time of molecules from similar molecular weight [38].

These results serve as promising preliminary data by suggesting a possible activation of osteocytes in old mouse bones. We then analyzed the effect of *in vivo* loading combined with Yoda2 injection in old mouse bone adaptation.

### 3.2. MicroCT scans analysis

Using microCT scans, we analyzed cortical bone parameters along the bone length for the different experimental conditions (Figure 2). We observed an increase in Ct.Ar, Ct.Th and Imin with loading and Yoda2. Paired SnPM1D analysis showed significant differences between groups Control+Vehicle compared to Load+Vehicle between 10% and 15% of the bone length, indicating that loading caused bone adaptation in specific regions, consistent with previous studies [17,29,35].The group Control+Yoda2 was significantly different from Control+Vehicle between 30% and 40% of the tibia length. When combining both factors, loading, and Yoda2 injection, we measured significant differences with the Control+Vehicle group between 10-15%, at 25%, and between 30-40% of the bone length.

**Figure 2:**
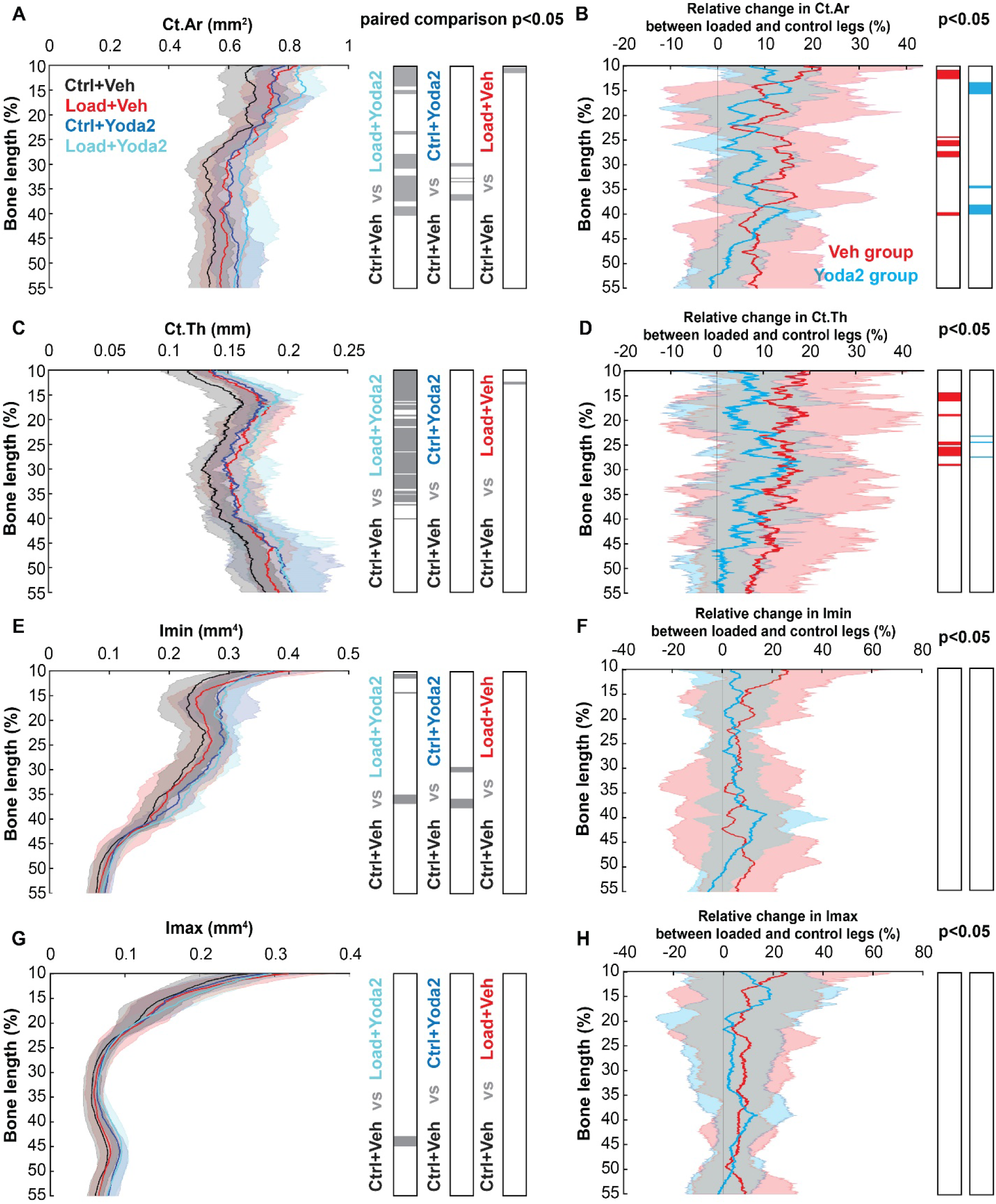
Cortical bone parameters analysis along the bone length. Cortical area along the bone length and paired statistical analysis of the different experimental conditions using SnPM1D (α=0.05). The grey regions of the bar graphs show regions of significant difference between the two tested groups. B) Relative changes in the cortical area between loaded and control legs of the vehicle and Yoda2 groups along the bone length. SnPM1D was used to test if the relative changes of the different groups were significantly different from 0 for α=0.05. Colored regions of the bar graphs show regions of significant relative changes. A similar approach was used for the other parameters C) Cortical thickness and D) Relative changes in cortical thickness. E) Minimum moment of area. F) Relative changes in the minimum moment of area. G) Maximum moment of area. H) Relative changes in the maximum moment of area. (Control + Vehicle: n=10 bones; Control + Yoda2 : n=10 bones; Loaded + Vehicle : n=8 bones; Loaded + Yoda2 : n=8 bones).

Quantification of the relative change in cortical bone parameters in the Vehicle and Yoda2 groups show the effect of loading on the bone parameters along the bone length (Figure 2.B.D.F.H). A positive relative change suggests that the loaded tibia has a larger cortical bone parameter than the contralateral leg of the same mouse. A relative change in Ct.Ar of about 10% for the Yoda2 group and 15% for the Vehicle group was measured along the bone length (Figure 2.B). Similar results were found for Ct.Th measurement (Figure 2.D). Significant relative changes in Ct.Ar and Ct.Th, in the loaded versus control legs were measured in both Vehicle and Yoda2 groups. In the Yoda2 group, relative changes in Ct.Ar were found at about 15% and between 35 and 40% of the bone length. Similar results were found in the Vehicle group, with the addition of significant relative change at about 25% of the bone length. No relative changes in Imin and Imax were measured in either group. However, paired SnPM1D analysis showed significant differences in Imin between the Control+Vehicle group versus the Control+Yoda2, and Load+Yoda2 groups at about 35% of the bone length, indicating that treatment with Yoda2 or Load+Yoda2 increased bone in the diaphysis. A significant difference was also found in Imax at about 45% of the bone length between Control+Vehicle and Load+Yoda2. These results suggest that although loading does not affect Imin and Imax with this loading protocol in aged mice, Yoda2 injections do influence those parameters.

We performed an analysis of cortical bone parameters in location-specific cross-sections and statistically tested significant differences using unbalanced 2-way ANOVA (Figure 3). The significant effect of loading was reported for Ct.Ar measurements at 25% of the bone length (p<0.05) and for Ct.Th at 25% (p<0.01), 37% and 50% (p<0.05) of the tibia length. On the other hand, the effect of Yoda2 injection was mostly reported for Imin measurements at 15%, 25%, and 37% (p<0.01) and for Ct.Ar at 50% of the bone length (p<0.05). We measured significant effects of both factors, loading and Yoda2 injection, on the Ct.Ar at 15% and 37% of the bone length (p<0.05), but also on Ct.Th at 15% of the bone length (p<0.01). Despite significant effects of both loading and Yoda2 at some locations, no interactions between the two factors were found according to 2-way ANOVA analysis. This result suggests an additive effect of the two factors (the effect of loading and Yoda2 add up) rather than a synergistic effect (factors combine to create a new effect greater than the sum of both factors).

**Figure 3:**
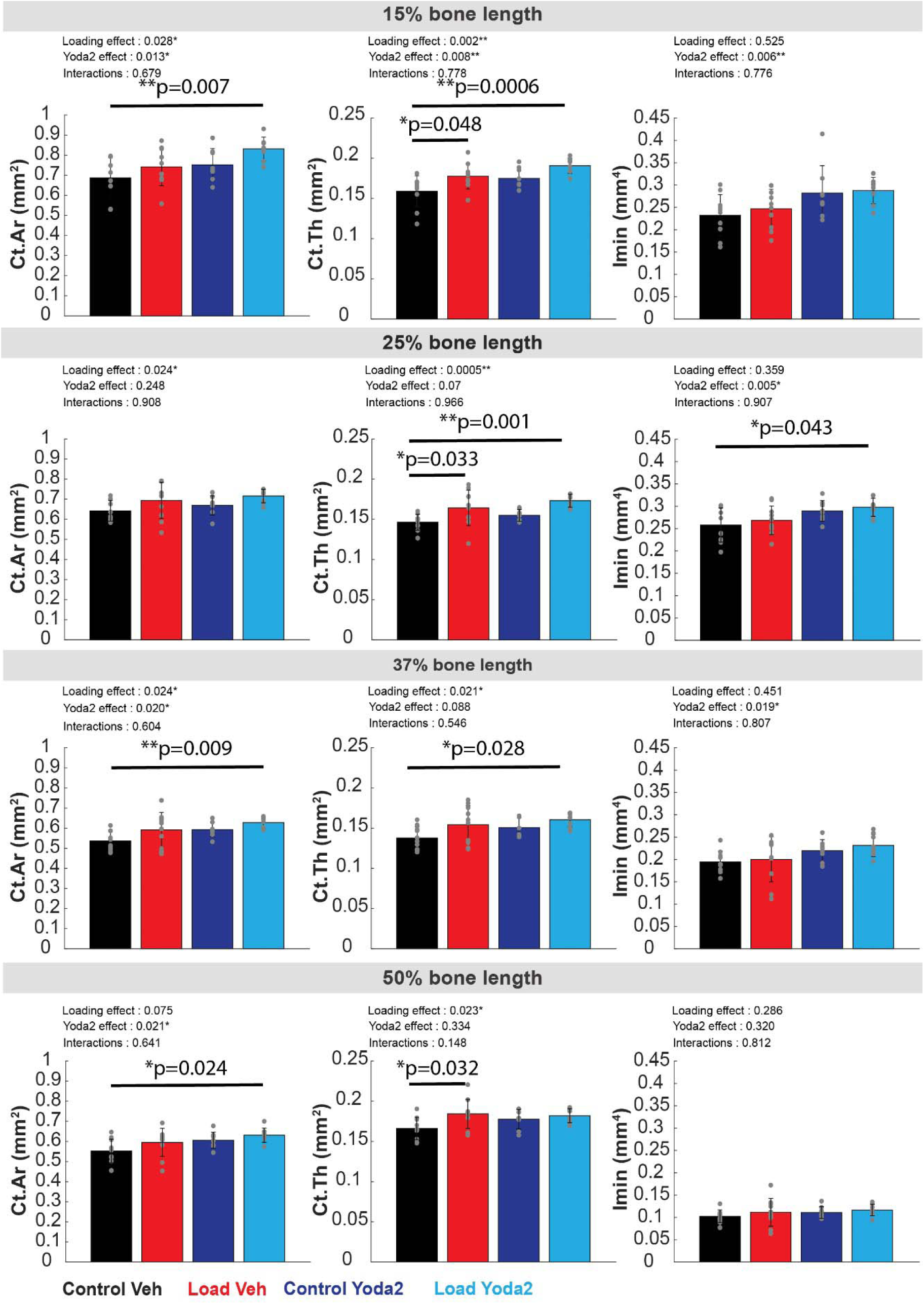
Cortical bone parameters analysis at specific locations of the bone. Ct.Ar, Ct.Th, and Imin were analyzed at 15, 25, 37, and 50% of the bone length. For each location, a 500 µm-thick section was considered for analysis. Unbalanced 2-way ANOVA was run to test for the significance effect of loading, Yoda2 injection, and interactions of both factors on bone parameters. P-values of individual effects and interactions were reported on the top left of each graph. A p-value inferior to 0.05 suggests a significant effect of the factor of interest at this location. In addition, we performed post-hoc Tukey-Kramer analysis to determine significant differences between groups. The results of the post-hoc analysis were reported by a horizontal black line and the corresponding p-value on top. *: p<0.05. **: p<0.01. (Control + Vehicle: n=10 bones; Control + Yoda2 : n=10 bones; Loaded + Vehicle : n=8 bones; Loaded + Yoda2 : n=8 bones).

Ma.Ar and Tt.Ar were not signicantly affected by loading nor Yoda2 injections (Figure 4). However the cortical bone fraction (Ct.Ar/Tt.Ar) was significantly greater in the Load+Yoda2 group compared to Control+Veh group at 15% of the bone length. A significant effect of loading and Yoda2 was found at 15% of the bone length according to the unbalanced two-way ANOVA. In addition, a significant effect of loading, but not of Yoda2, was found at 37% of the bone length (Figure 4).

**Figure 4:**
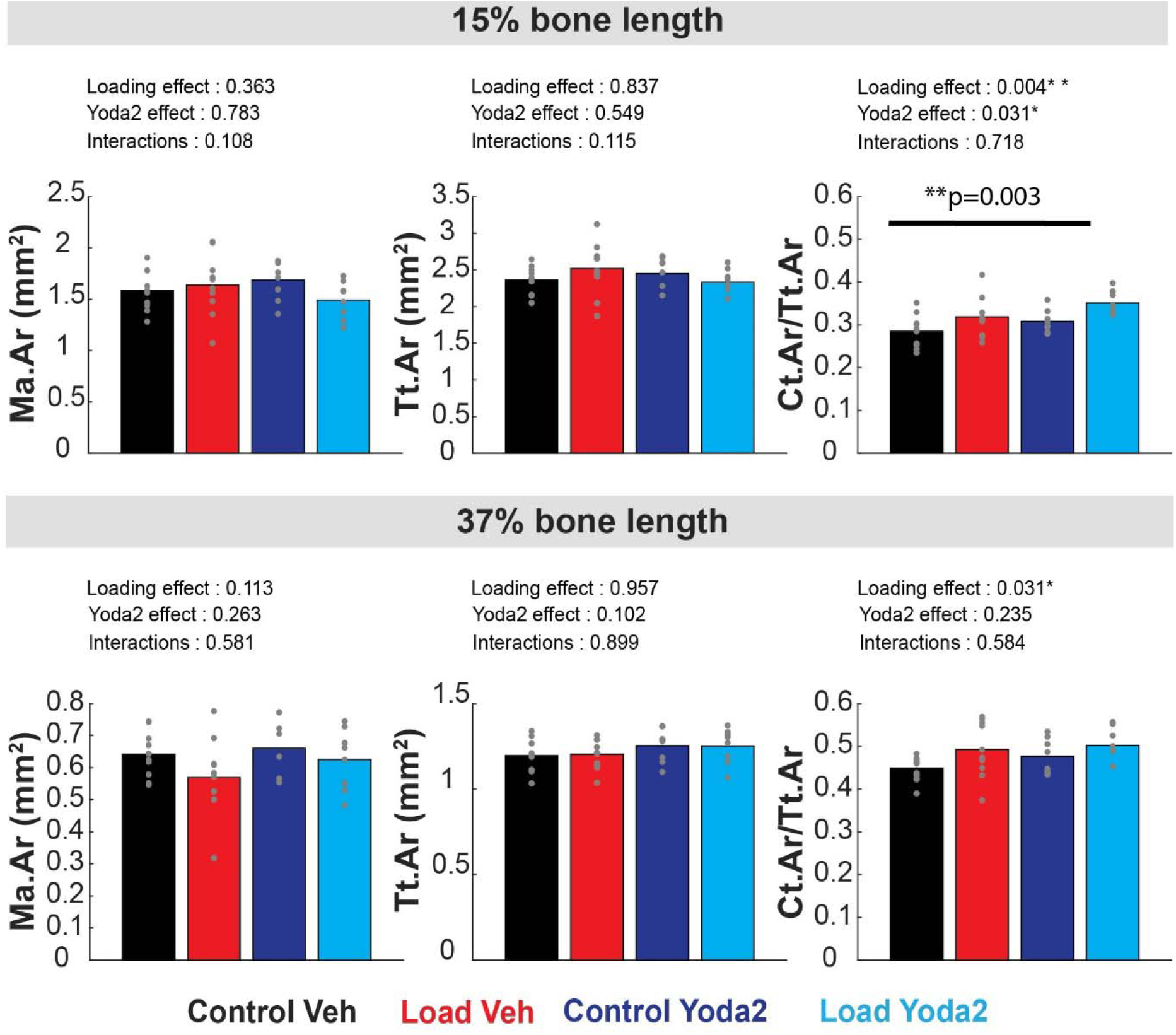
Marrow area, total area, and cortical bone fraction measured at 15% and 37% of the bone length. For each location, a 400 µm-thick section was considered for analysis. Unbalanced 2-way ANOVA was run to test for the significant effect of loading, Yoda2 injection, and interactions of both factors on bone parameters. P-values of individual effects and interactions were reported on the top left of each graph. A p-value inferior to 0.05 suggests a significant effect of the factor of interest at this location. In addition, we performed post-hoc Tukey-Kramer analysis to determine significant differences between groups. The results of the post-hoc analysis were reported by a horizontal black line and the corresponding p-value on top. *: p<0.05. **: p<0.01. (Control + Vehicle: n=10 bones; Control + Yoda2 : n=10 bones; Loaded + Vehicle : n=8 bones; Loaded + Yoda2 : n=8 bones).

We quantified the gross cortical porosity (1-(Exp Ct.Ar/Theo Ct.Ar)). Results showed a significant decrease in porosity with Yoda2 injections at 15% and 37% of the bone length (Figure 5).

**Figure 5:**
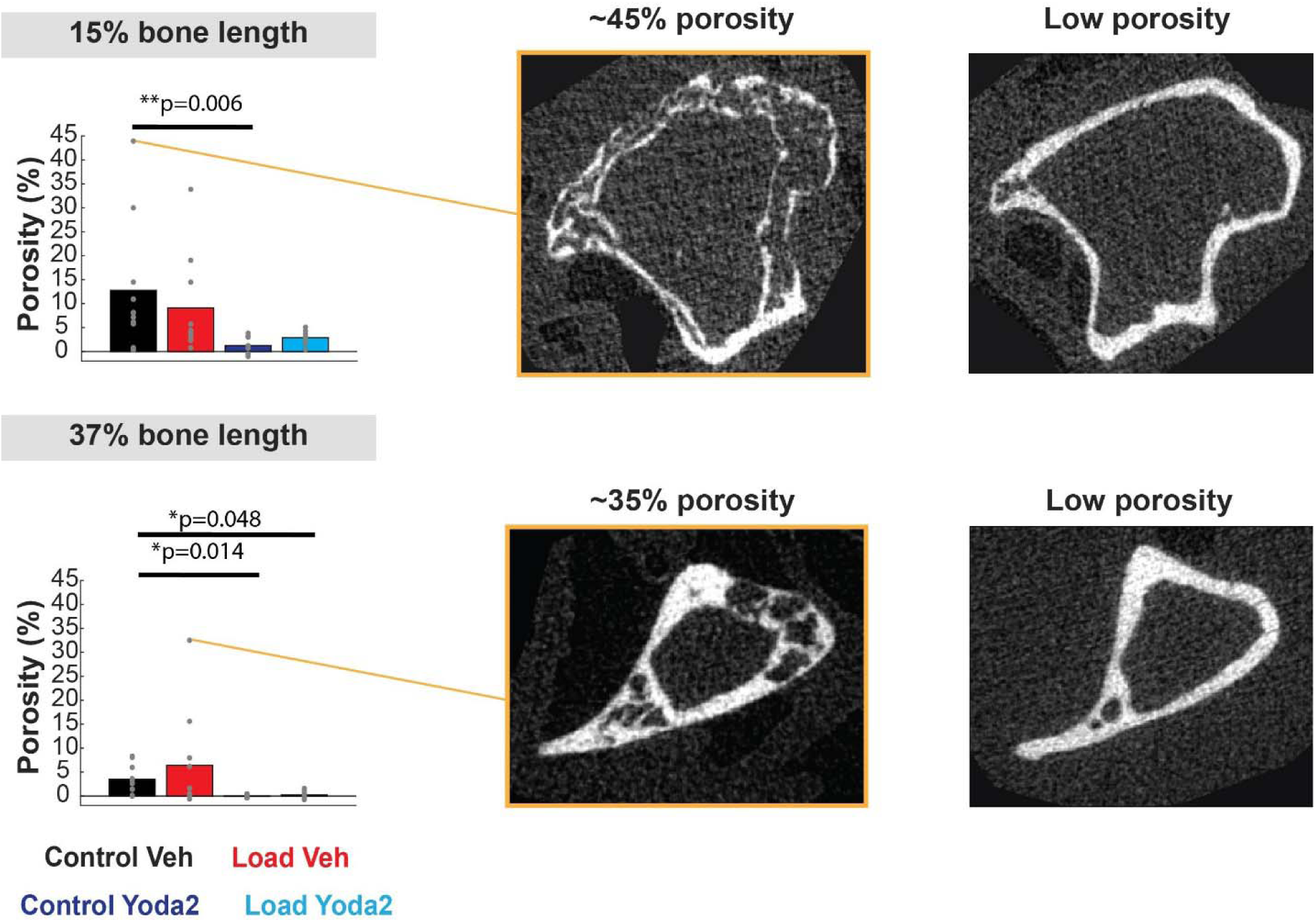
Cortical porosity measurement at 15% and 37% of the bone length. For each location, a 400 µm-thick section was considered for analysis. Examples of bone cross-section with high and low porosity are presented. Kruskal–Wallis test followed by post-hoc comparison were performed to test for significant differences between the groups.

We also measured trabecular bone parameters and noticed very few trabeculae in the region analyzed, which is characteristic of old mice bones. Trabecular bone analysis showed an effect of loading on trabecular thickness but not on the other parameters (Figure 6). Yoda2 injection did not show any significant effect on trabecular bone parameters according to an unbalanced 2-way ANOVA analysis.

**Figure 6:**
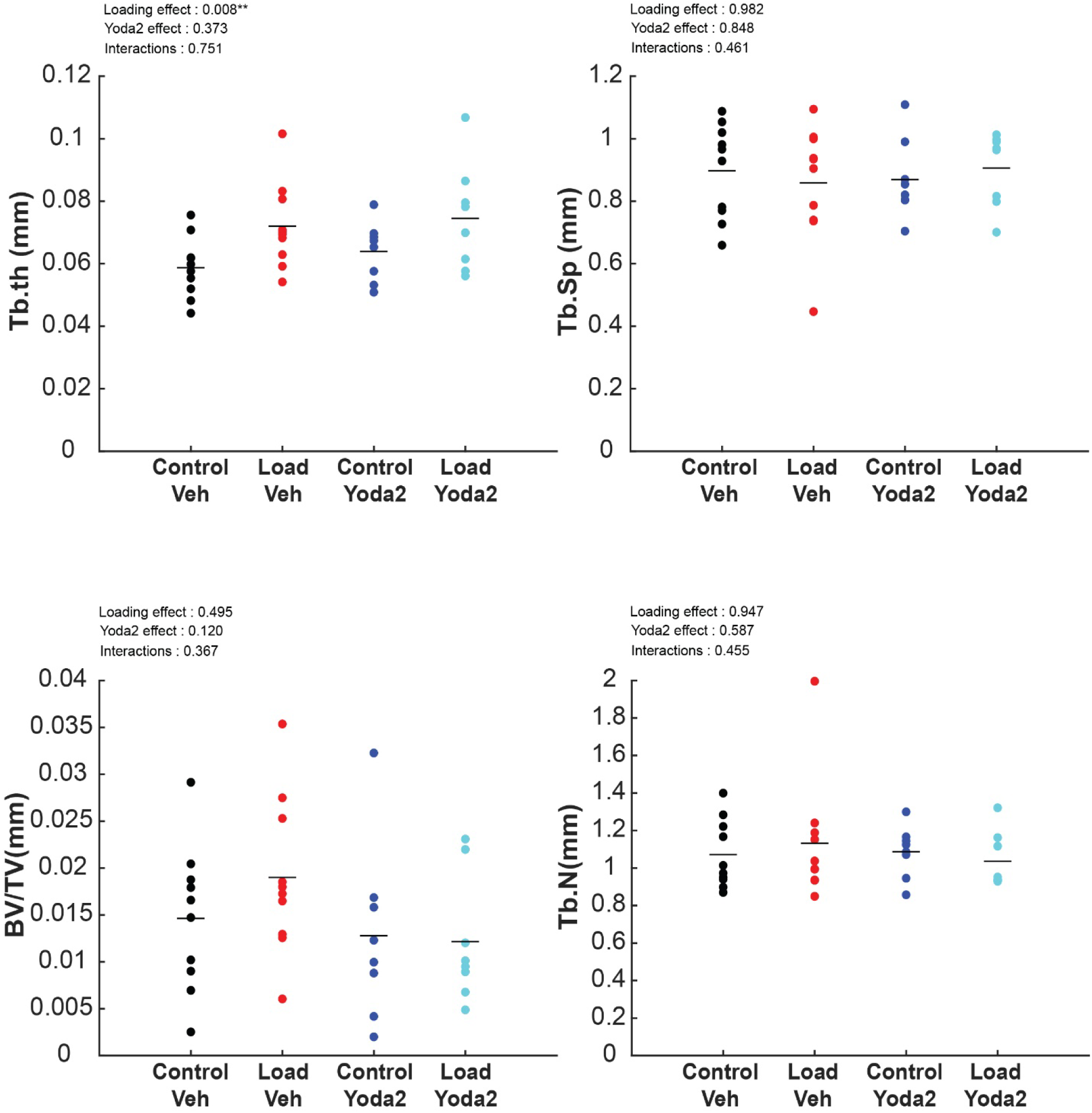
Trabecular bone parameters analysis. Unbalanced 2-way ANOVA was run to test for the significant effect of loading, Yoda2 injection, and interactions of both factors on trabecular bone parameters. P-values of individual effects and interactions were reported on the top left of each graph. A p-value inferior to 0.05 suggests a significant effect of the factor of interest at this location. *: p<0.05. **: p<0.01. In addition, post-hoc Turkey Kramer tests were performed. (Control + Vehicle: n=10 bones; Control + Yoda2 : n=10 bones; Loaded + Vehicle : n=8 bones; Loaded + Yoda2 : n=8 bones).

To summarize, 2 weeks of loading and Yoda2 injections were effective at increasing cortical bone parameters in 22-month-old mice compared to non-loaded tibia with vehicle injection. Yoda2 and loading seem to have significant effects on cortical bone parameters at various locations along the bone length. Statistical analysis suggests an additive effect of loading and Yoda2 injection on cortical bone parameters rather than synergetic effects.

## 4. Discussion

In this work, we investigated the effect of 2 weeks of tibia loading combined with injections of Yoda2 on the bone mechnaoadaptation response of 22-month-old mice. First, we used a transgenic mouse model and *in vivo* two-photon microscopy to assess the potential of Yoda2 to induce osteocyte activation in old mouse bones. Then, we stimulated old mice bones with tibia loading and Yoda2 injections for 2 weeks. We found that mechanical and chemical stimulation results in an increase in cortical bone parameters. Statistical analysis suggests an additive effect of the two stimuli.

Our calcium signaling experiment hints at an increase in the percentage of responding osteocytes 1h after Yoda2 injection *in vivo* compared to baseline. The percentage of cells responding was found to be back to baseline about 2h after injections. Calcium signaling influxes attributed to the Yoda2 injection were observed to last on average about 40 seconds. This result is consistent with previous calcium signaling time characteristics reported *in vitro* and *ex vivo* following mechanical stimulation [12,37,39,40]. However, these time characteristics differ from recent *in vivo* measurements following bending of the mouse metatarsal [41,42]. Further investigation of the time characteristic of osteocytes calcium signaling *in vivo* is necessary to explore potential differences between chemically activated and mechanically activated osteocytes calcium signaling.

A major limitation of our calcium signaling investigation is the number of replicates. We performed baseline measurements in two transgenic mice, imaging 2 different ROIs per mouse for a total of 180 cells analyzed. The effect of Yoda2 was measured in one transgenic mouse, at four different time points for a total of about 45 cells analyzed per time point. A second mouse, used for baseline measurement, could not be used to analyze the effect of Yoda2 injection due to experimental challenges. However, we used the results of the one mouse as preliminary data that were encouraging to perform *in vivo* Yoda2 injections in old mice with the objective of stimulating *in vivo* osteocytes.

Osteocyte mechanosensitivity has been commonly studied *in vitro* via calcium signaling measurements in response to mechanical stimuli [39,40,43]. In addition, *Ex vivo* tibia loading was used to characterize the effect of mechanical loading on osteocytes calcium signaling within the tissue [37] and showed an increase in the percentage of responding osteocytes with the increase of load-induced strain magnitude Calcium signaling influx duration was captured to last between 30 sec to 1 min long, which is consistent with *in vitro* calcium influx time characteristics. Using the same method, Morrell et al. showed that the percentage of cells responding to loading decreases with age[12]. *Ex vivo* models preserve the structural environment of the cells which addresses some of the major limitations of *in vitro* models. However, the lack of pressure due to the extraction of the sample may interfere with the mechanical signal sensed by the osteocytes. Recently, osteocyte calcium signaling response to mechanical loading has been investigated *in vivo*. [26]. They used a transgenic mice model with a GCaMP reported expressed in osteocytes driven by *Dmp1* expression. The authors reported that the percentage of responding osteocytes increased with loading magnitude. Interestingly, time characteristics of calcium influx, measured *in vivo* were reported to be in line with the loading frequency, lasting about 1 second. This duration of calcium influx does not match previous results reported *in vitro* and *ex vivo*.

Investigating calcium signaling in osteocytes from *in vitro* to *in vivo* studies inform on the osteocyte’s activation in response to loading and potential factors that could affect the cells mechanosensitivity, such as aging. Discrepancies in the literature remain regarding the time characteristics of calcium signaling *in vivo* compared to ex-vivo and *in vitro* models. In this work, we performed *in vivo* calcium signaling imaging of osteocytes after injection of a Piezo1 agonist to hint its efficacy to trigger old bone cells activation *in vivo*. Inducing calcium signaling in osteocytes could participate in triggering a bone adaptation response in samples that typically do not respond well to only mechanical loading. Preliminary data suggest an increase in osteocytes responding to Yoda2 injection *in vivo*, and the calcium signaling influx presents time characteristics similar to what has been reported *ex-vivo* and *in vitro* experiments.

To mechanically load the mice, we used a 7N trapezoid loading profile that we found to be efficient at inducing bone formation in adult bone [29]. This loading profile was designed to induce high fluid flow velocity in adult mouse tibiae and lower strain magnitude than the commonly used loading profiles. In the presented study, we observed that this new loading profile enabled bone adaptation in old mouse bone and had a significant effect on multiple cortical bone parameters and trabecular bone thickness. Previous studies investigating old mouse bone response to loading often reported a lack of adaptation [44]. Javaheri et al suggested that increasing the load magnitude for a couple loading bouts can help prime bone adaptation in old mice [17]. Here we reported that a low load magnitude loading profile can stimulate old mouse bone adaptation, most likely due to the high load rate. This high load rate was found to induce high fluid low velocity in simulated mouse tibia loading [29]. Thus, mechanical loading alone seems able to trigger old mouse bone adaptation. However, careful design of the loading for further optimization is required.

We found that Yoda2 injections had a significant effect on cortical bone parameters such as minimum moment of area and cortical area. To our knowledge, this is the first evidence demonstrating the effects of Yoda2 injections *in vivo* in mice. Our results suggest that Yoda2 injection on its own can help promote cortical bone parameters. These results are consistent with previous studies investigating the effect of Yoda1 injection, another Piezo1 agonist, in adult mice [20,23]. Li et al. [20] found that 2 weeks of Yoda1 injections increased cortical thickness and trabecular bone volume fraction, which differ from our findings. Indeed, we did not observe any effects of Yoda2 injections on trabecular bone parameters. This lack of effect may be attributed to the very low number of trabeculae present in the 22-month-old mice used in this study. It is likely that a 2-week period of Yoda2 injections was insufficient to induce measurable benefits in the nearly absent trabecular compartment. A longer injection period might be necessary, or it might not be possible to rebuild trabecular bone after it has been lost. A previous study investigated the effect of one month of Yoda1 injection in 20-month-old mouse trabecular bone and found an increase in trabecular bone parameters in treated mice compared to untreated old mice [23].

Combining Yoda2 injections and tibia loading resulted in the increase in cortical bone parameters compared to the control group. The statistical analysis suggested an additive effect of loading and Yoda2 injections on the measured bone parameters. We observed beneficial effects of this combination after two weeks of treatment only which is a promising result for future treatment strategy development for age-related bone mass loss. A recent study showed that a month of moderate tibia loading and Yoda1 injection was efficient at increasing cortical and trabecular bone parameters in mature mice (50 weeks old) whereas moderate loading itself did not have a significant effect on bone parameters [24]. Similar to our study, this previous study also found an additive effect of loading and Yoda1 injections in mature mouse bone. The researchers used mature mice, which are prone to losing bone mass over time but are expected to still possess trabecular bone. In contrast, our study examined the combined effects of a new loading protocol and the new Piezo1 agonist Yoda2 over 2 weeks in 22-month-old mice. These older mice had very few trabeculae remaining.

## 5. Conclusion

In this study, we reported an increase in cortical bone parameters in response to tibia loading combined with Yoda2 injection, a new Piezo1 agonist, in 22-months-old mouse bones. An additive effect was found between the two stimuli in cortical bone parameters. This work shows that the combination of mechanical and chemical stimulation presents a promising strategy to promote bone adaptation and restore bone mass loss issues in the elderly.

## Acknowledgments

This work was funded by the National Science Foundation CMMI # 2010010

We thank the Institute for Chemical Imaging of Living Systems (RRID:SCR_022681) at Northeastern University for consultation and instrument support.

## Author Contributions

**Quentin A Meslier:** Conceptualization, Methodology, Investigation, Formal analysis, Visualization, Writing – original draft, Writing – review & editing. **Robert Oehrlein:** Formal analysis. **Sandra J. Shefelbine**: Conceptualization, methodology, Writing – review & editing, Supervision, Funding acquisition.

