## Supplemental methods for "Combined effects of mechanical loading and Piezo1 chemical activation on 22-months-old female mouse bone adaptation"

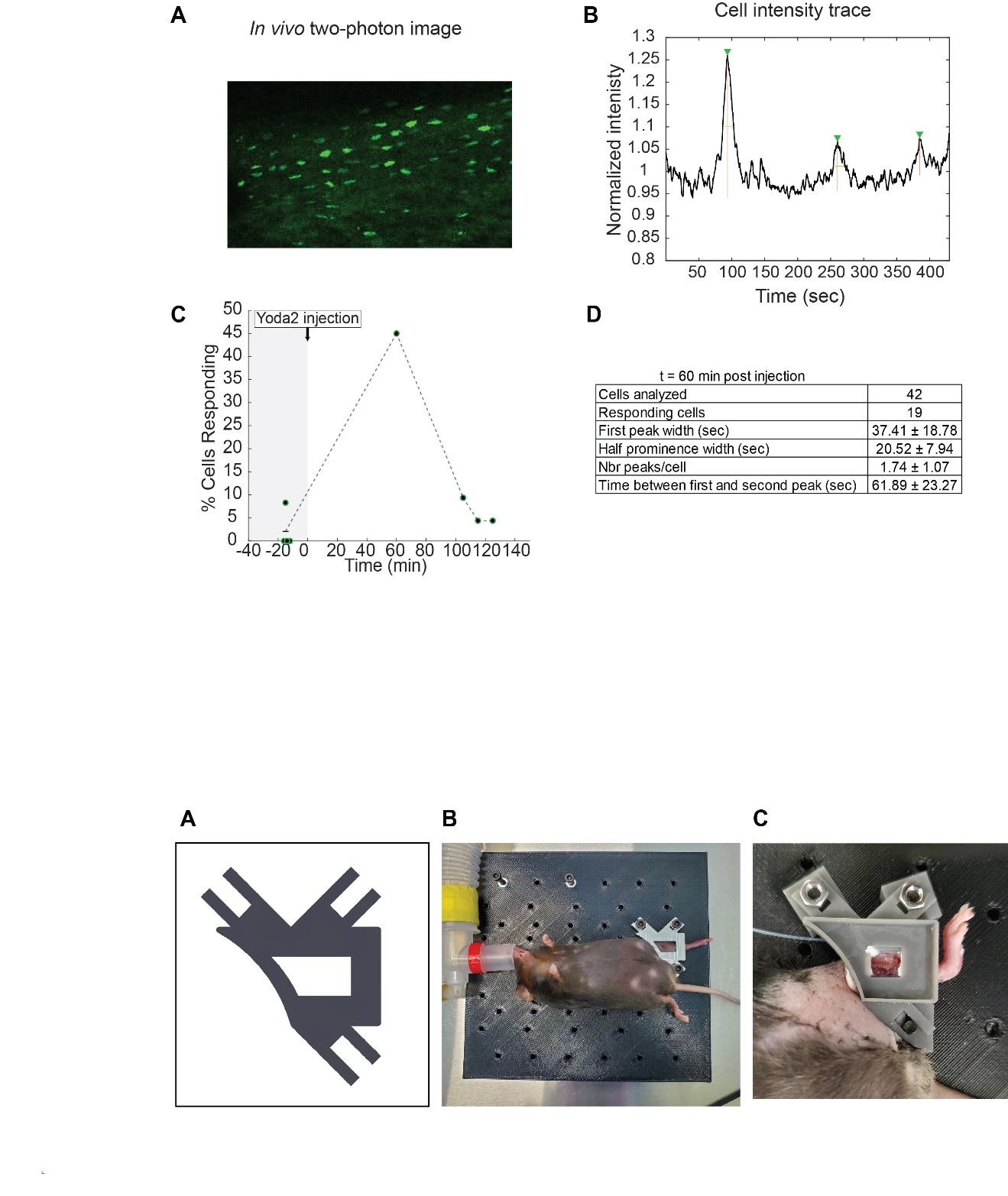

Supplemental Figure 1: Tibial plate for *in vivo* imaging.

A) CAD file of the tibial plate. B) Example of mouse tibia fixation using the tibial plate and the custom platform. C) Preparation of the mouse tibia for imaging. The tibia is surgically exposed and glued to the tibia plate. Sealant is applied at the plate-tibia interface to avoid leakage of the immersion solution. The plate is screwed to the custom platform for immobilization of the tibia.

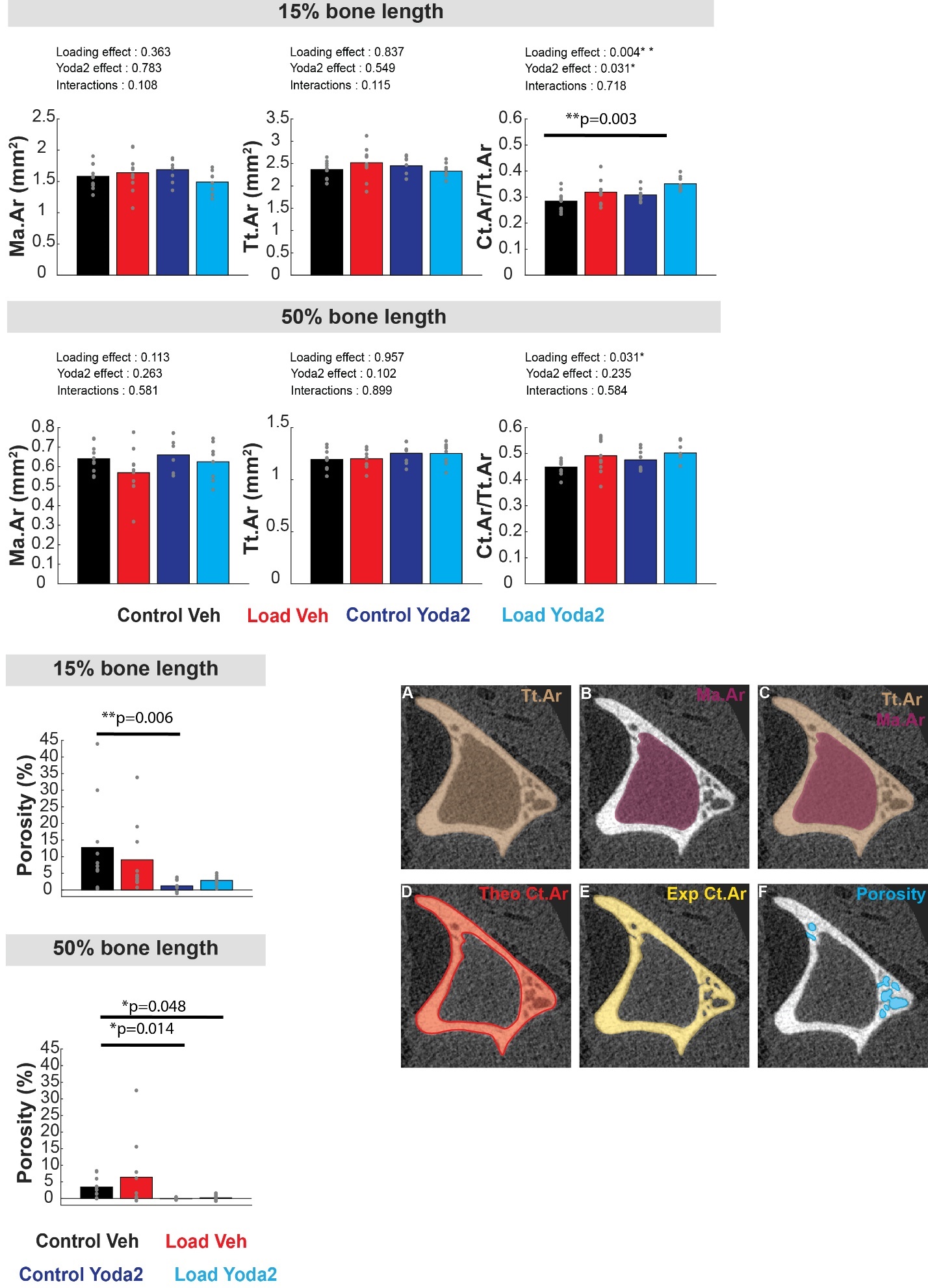

Supplemental Figure 2: Illustration of the method used for calculation of the cortical gross porosity in old mouse tibia midshaft. A) Total area (Tt.Ar) was defined by the cortical bone's outer surface. B) The inner surface of the cortical bone defined marrow area (Ma.Ar). C)-D) Theoretical cortical area corresponding to the subtraction of the total area by the marrow area. E) Experimental cortical area was measured using BoneJ in ImageJ. F) Gross cortical porosity. This parameters was reported as a percentage via the following formula: 1- (Experimental Ct.Ar/Theoretical Ct.Ar).
